# Temporal ordering of input modulates connectivity formation in a developmental neuronal network model of the cortex

**DOI:** 10.1101/355800

**Authors:** Caroline Hartley, Simon Farmer, Luc Berthouze

## Abstract

Preterm infant brain activity is discontinuous; bursts of activity recorded using EEG (electroencephalography), thought to be driven by subcortical regions, display scale free properties and exhibit a complex temporal ordering known as long-range temporal correlations (LRTCs). During brain development, activity-dependent mechanisms are essential for synaptic connectivity formation, and abolishing burst activity in animal models leads to weak disorganised synaptic connectivity. Moreover, synaptic pruning shares similar mechanisms to spike-timing dependent plasticity (STDP), suggesting that the timing of activity may play a critical role in connectivity formation. We investigated, in a computational model of leaky integrate-and-fire neurones, whether the temporal ordering of burst activity within an external driving input could modulate connectivity formation in the network. Connectivity evolved across the course of simulations using an approach analogous to STDP, from networks with initial random connectivity. Small-world connectivity and hub neurones emerged in the network structure - characteristic properties of mature brain networks. Notably, driving the network with an external input which exhibited LRTCs in the temporal ordering of burst activity facilitated the emergence of these network properties, increasing the speed with which they emerged compared with when the network was driven by the same input with the bursts randomly ordered in time. Moreover, the emergence of small-world properties was dependent on the strength of the LRTCs. These results suggest that the temporal ordering of burst activity could play an important role in synaptic connectivity formation and the emergence of small-world topology in the developing brain.

## Introduction

Network connectivity shapes activity and modulates information transfer in the brain. For example, small-world network architecture allows efficient integration and segregation of information [1], and hub neurones or regions play a key role in carrying information throughout the brain [2]. This structured connectivity emerges during early human brain development; a small-world modular network organisation with hub nodes can be observed in preterm diffusion and functional MRI, with a significant increase in small-world topology between 30 and 40 weeks’ gestation [3, 4].

The major period of connectivity formation and refinement in the cortex starts during foetal development from approximately 20 weeks’ gestation and continues for the first few years of postnatal life [5,6]. MEG recordings of foetal brain activity and EEG recordings from preterm infants are characterised by discontinuous activity - bursts of slow wave oscillations with nested high frequency activity are interspersed within periods of apparent electrical silence [7]. These bursts can occur in response to sensory stimulation [8–10], or following movement [11], but the majority occur spontaneously in the background EEG [11]. Spontaneous bursts are thought to originate from regions such as the subplate [12, 13], a transient population of neurones present in early development [14], and may also relate to activity in the insula [15]. Neuronal activity is crucial for connectivity formation [16], and blocking or reducing burst activity during critical developmental periods, for example, through removal of the subplate in animal models, leads to abnormal cortical network connectivity, with weak thalamocortical connectivity [17] and loss of cortical columnar structure [18, 19].

The temporal organisation of neuronal activity in early development may also play a key role in connectivity formation - rearing fish in an environment with stroboscopic illumination prevents the refinement of retinotectal maps [20] and periodic electrical stimulation of the ferret optic nerve results in altered orientation selectivity in the cortex [21]. Moreover, activity-dependent mechanisms at the synaptic level are key to connectivity refinement in the developing brain, and synaptic pruning shares similar molecular pathways with long-term depression (LTD) in the adult brain [22], suggesting that the temporal organisation of activity may play a critical role in connectivity formation. Recently it has been demonstrated that the burst activity in preterm EEG exhibits scale-free properties [23], and that the bursts do not occur randomly in time but follow a complex temporal ordering, known as long-range temporal correlations (LRTCs) [24]. Thus, in the preterm neonatal brain, the timing of any given EEG burst is correlated with the time of occurrence of all previous bursts of EEG activity [24]. Whether this correlated temporal structure of the timing of burst activity affects connectivity formation in the developing brain is an important open question.

Here we consider this question by investigating connectivity formation in a simple activity-dependent neuronal network model of the cortex. Motivated by the excitatory role of GABA within the developing brain [25], we consider a model where all connections are excitatory. We assume the cortex is driven by bursts of activity from a non-cortical source such as the subplate [12, 13]. We compare connectivity formation in a network driven by bursts which exhibit LRTCs, to the network connectivity that emerges when the network is driven by bursts with random temporal ordering. This preserves the inter-burst interval distribution whilst changing the temporal correlations in the ordering of the inter-burst intervals. We also compare with the connectivity in networks driven by periodic bursts. Finally, we investigate the relationship between network connectivity parameters and the strength of LRTCs within the burst temporal organisation. We test the hypothesis that LRTCs in the burst activity of the external input promotes the emergence of small-world connectivity in the developing cortex.

## Results

We investigated connectivity formation in directed networks of leaky integrate-and-fire neurones. Whilst this work is motivated by the observation of LRTCs in the EEG of preterm infants, i.e. an observation at the macroscopic scale, LRTCs have been demonstrated at multiple levels of the nervous system in adults, including in the spontaneous spiking of individual neurones [26]. Although the external input which provides the LRTCs within our model is on the macroscopic scale, with bursts originating from non-cortical sources such as the subplate [12, 13], our focus here is on the effect such macroscopic phenomenon has on connectivity formation at the microscopic (neuronal) scale.

Individuals neurones in our model received both this external input and input from connected neurones when these neurones fired. To ensure that neuronal firing did not saturate the network or die out completely, synaptic weights were updated according to the number of connections within the network (see *Methods*), which can be considered a form of homeostatic plasticity [27]. The external input to the network was a bursty input, Fig. 1A. In all cases the bursts themselves were of a fixed duration and amplitude; differences in the driving input were only reflected in the temporal ordering of the bursts, i.e, in the ordering of the interburst intervals (IBIs) - the time between the bursts. In the case where the network was driven with burst dynamics that exhibit LRTCs, the sequence of IBIs exhibited long-range temporal correlations with a Hurst exponent (*H*) greater than 0.5. In the shuffled input case, the same IBIs were randomly re-ordered in time, giving a Hurst exponent of the sequence of IBIs of *H* ≈ 0.5. Thus, in both cases the external input had the same IBI distribution, but the temporal ordering of the bursts was changed and in the latter case the temporal correlations were lost.

**Figure 1:**
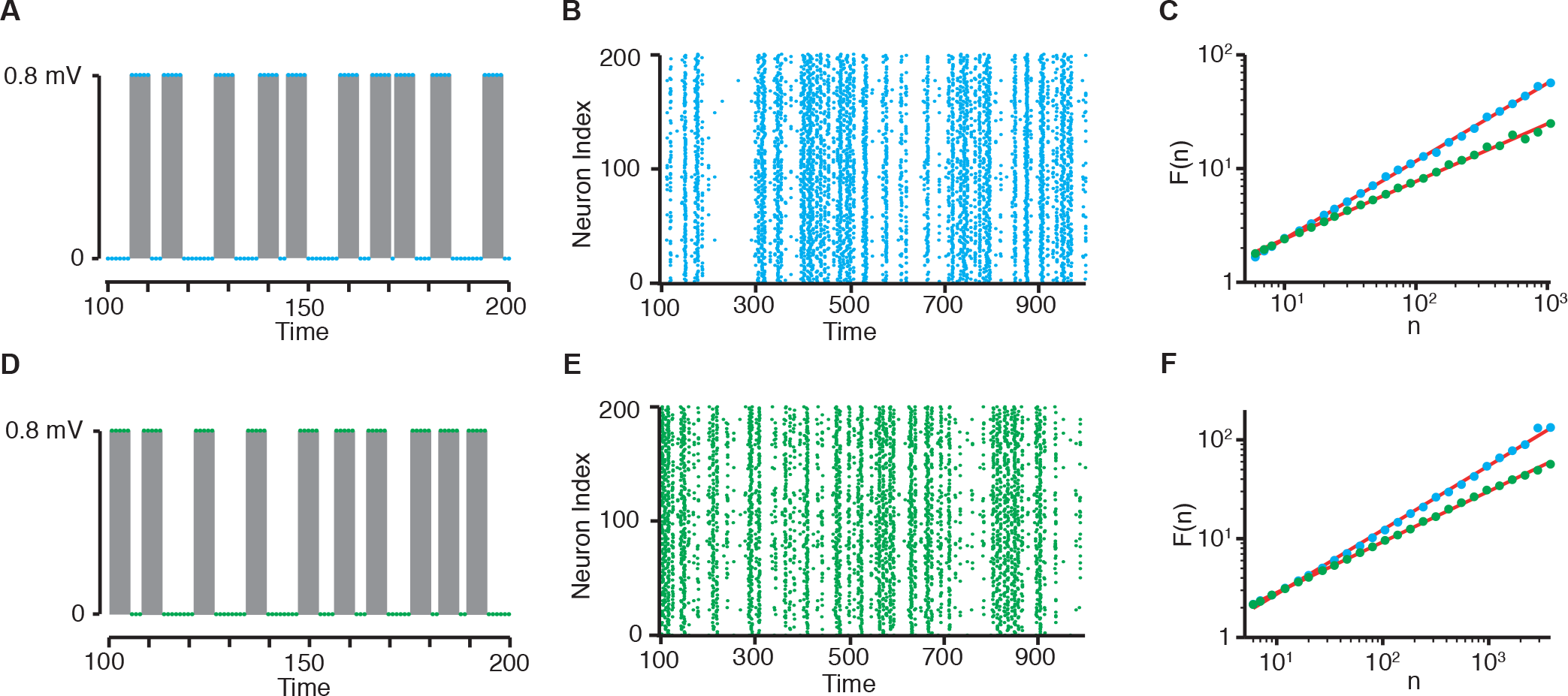
Network input and burst dynamics. (A,D) Networks were driven by a bursty input. Bursts of activity (grey shaded region) were of fixed duration and amplitude, and interspersed within periods of silence - inter-burst intervals (IBIs). (A) Example burst dynamics for the first few bursts within a simulation where IBIs exhibited LRTC, and (D) the same IBIs randomly shuffled. (B,E) Raster plot of network firing at the start of a simulation, demonstrating that burst dynamics also occur within the network. Firing dynamics are shown for the network driven with LRTCs (B, blue), and driven with the same IBIs randomly shuffled (E, green). (C) DFA plots of the IBI sequences for the network external input across simulations of length 100, 000. The Hurst exponent is estimated by the slope of the line of best fit, which in these examples were *H* = 0.68 (with LRTC input, blue), and *H* = 0.51 (with the same input shuffled in time, green). (F) DFA plots for the burst dynamics of the network firing across the same simulations. For these examples the Hurst exponents of the network firing were estimated as *H* = 0.65 when the network was driven by an external input which exhibited LRTCs (blue), and *H* = 0.51 when the network was driven with the same input randomly re-ordered in time (green). *n* is the box size and *F*(*n*) is the root mean square of the detrended signal across the box (see *Estimation of the Hurst exponent in the Methods*).

Driving the neuronal networks with bursty input also led to bursts within the network (Fig. 1B,E). The Hurst exponent for the sequence of IBIs was determined using detrended fluctuation analysis (DFA). The DFA exponents for the sequences of IBIs in the network firing dynamics reflected those of the corresponding DFA exponents of the external input (Fig. 1C,F), indicating that the temporal characteristics of the driving input were transmitted to the network activity itself.

### Emergence of small-world topology and hub neurones

Network connectivity was allowed to evolve during simulations using an approach analogous to spike-timing dependent plasticity (STDP). The likelihood of gaining a connection between any two neurones was increased if the presynaptic neurone frequently fired just before the postsynaptic neurone, whereas the likelihood of losing a connection was increased if the postsynaptic neurone frequently fired before the presynaptic neurone. Connections were then lost or gained if this likelihood measure reached set thresholds for gaining and losing connections (see Methods). Network connectivity was initially random, with 40% connectivity in a network with 200 neurones (alterations to these parameters are considered in the section *Varying network size and density*). As with STDP in the adult brain [28], and as similar mechanisms to LTD play a dominate role in neuronal network development [22], we initially set depression to be slightly stronger than potentiation. We also modelled alternatives, i.e. equal amounts of depression and potentiation and stronger potentiation than depression, which are described below (see section *Varying the plasticity parameters*).

With slightly stronger depression than potentiation, and driven with burst dynamics which exhibit LRTCs, the network on average lost connections across the course of the simulation and, although the effect size is relatively small, the normalised clustering coefficient and small-world index increased. Fig. 2 shows the mean proportion of connections (i.e. the number of connections divided by the number of all possible connections within the network), normalised clustering coefficient, normalised mean path length and small-world index from 20 simulations, which were all driven with external input which exhibited LRTC with *H* ≈ 0.7. At the end of the simulations, the node in degree distribution is skewed, with some neurones showing much higher degree than others, indicating the presence of hub neurones (Fig. 2D). Thus, across the course of the simulation a small-world topology and hub neurones emerge. Moreover, the speed at which the proportion of connections, normalised clustering coefficient and small-world index changes is high at the start of the simulations, indicating that the network rapidly evolves to have connectivity with small-world properties.

**Figure 2:**
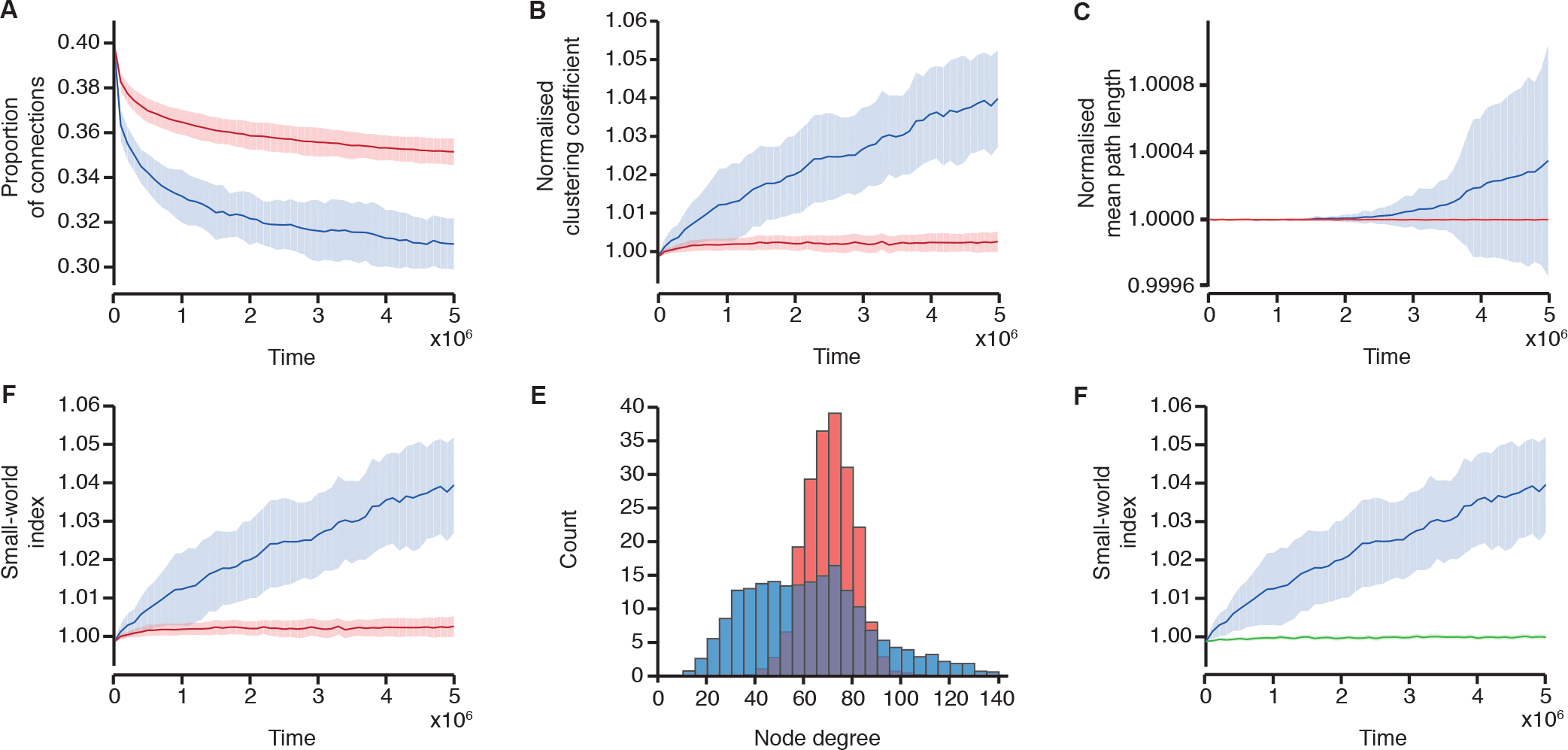
Emergence of small-world topology and hub neurones. Network parameters across the course of simulations in networks driven with burst activity which exhibits LRTCs in the IBIs (blue), with input with the same IBIs randomly shuffled in time (red) and with periodic bursts (green). Solid lines indicate the mean across 20 simulations, and the shaded areas indicate the standard deviation. (A) The proportion of connections in the network, (B) the normalised clustering coefficient, (C) the normalised mean path length, and (D) the small-world index across the course of the simulations in networks driven with burst activity which exhibits LRTCs in the IBIs (blue) and with input with the same IBIs randomly shuffled in time (red). (E) Average in degree distributions at the end of the simulations with the networks driven by burst activity which exhibits LRTCs in the IBIs (blue) and by the input with the same IBIs shuffled (red). (F) The small world index in networks driven with burst activity which exhibits LRTCs in the IBIs (blue) and with periodic bursts (green).

In contrast, when the network is driven with the same external input but in which the burst order has been randomly shuffled in time (and therefore does not exhibit LRTCs but instead *H* ≈ 0.5), the rate of change in the network parameters at the start of the simulation is much lower than with external input which exhibits LRTCs, Fig. 2. Thus, whilst over the course of the simulation on average connections are still lost, the final network has a higher proportion of connections. The normalised clustering coefficient and small-world index exhibit very little change compared with when the networks are driven by bursts which exhibit LRTCs. Moreover, the resultant degree distribution is approximately normal and does not indicate the presence of hub neurones, Fig. 2E.

Comparison of the speed at which the network properties evolve in the network driven by external input which exhibits LRTCs with the case where the external input is randomly shuffled in time suggests that there is an early window during which the network is highly sensitive to the temporal organisation of the external input. Whilst the level of activity within the external input across the whole simulation is identical in both cases, we also ascertained that this difference in the speed of emergence of small-world properties was not related to differences in the level of activity of the external input at the start of the simulation. The average rate of input at the start of the simulation was similar when the input exhibited LRTCs and when the IBIs were randomly shuffled in time (Fig. S1). Moreover, considering individual examples with similar levels of the external input at the start of the simulations, those networks driven with input which exhibits LRTCs have different trajectories of their network properties compared with those driven with shuffled input (Fig. S2). Therefore, the difference in the speed of emergence of the network properties between the networks driven with external input with LRTCs in the IBIs and the shuffled input is not because the level of input to the network is different at the start of the simulation.

We chose to compare the network driven by external input which exhibits LRTCs with the case where the external input is randomly shuffled in time as shuffling the inter-burst intervals preserves the other properties of input, namely the distribution of the IBIs, whilst destroying the temporal ordering. We also investigated what happened to the network when it was driven with a periodic external input - where all the IBIs were identical in length and set to the average duration of the IBIs in the LRTC and shuffled cases (see *Methods*). With periodic external input the rate of change in the network parameters is also much lower than when the external input has LRTCs, Fig. 2F, though it should be noted that as all IBIs are equal in length the periodic external input does not preserve the overall IBI distribution, unlike the case where the external input is randomly shuffled.

The difference in emergence of small-world properties when the external input exhibits LRTCs leads to the question as to whether the magnitude of the LRTCs also affects connectivity formation. Therefore, we next investigated how driving the network with burst activity with IBIs which exhibit different strengths of LRTCs alters the rate of emergence of small-world properties.

### Varying the Hurst exponent modulates connectivity formation

We investigated driving the network with burst dynamics which had 4 different levels of Hurst exponent: *H* ≈ 0.5, 0.6, 0.7 and 0.8. In all cases, on average the network lost connections, but networks driven by external input with higher Hurst exponents lost connections at a faster initial rate and quickly evolved to a network with small-world properties, see Fig. 3. The average Hurst exponent estimated using DFA in preterm infants is 0.68, with a range of 0.55 − 0.81 [24], so the levels of Hurst exponent seen in preterm infants fall within the range of data simulated here.

**Figure 3:**
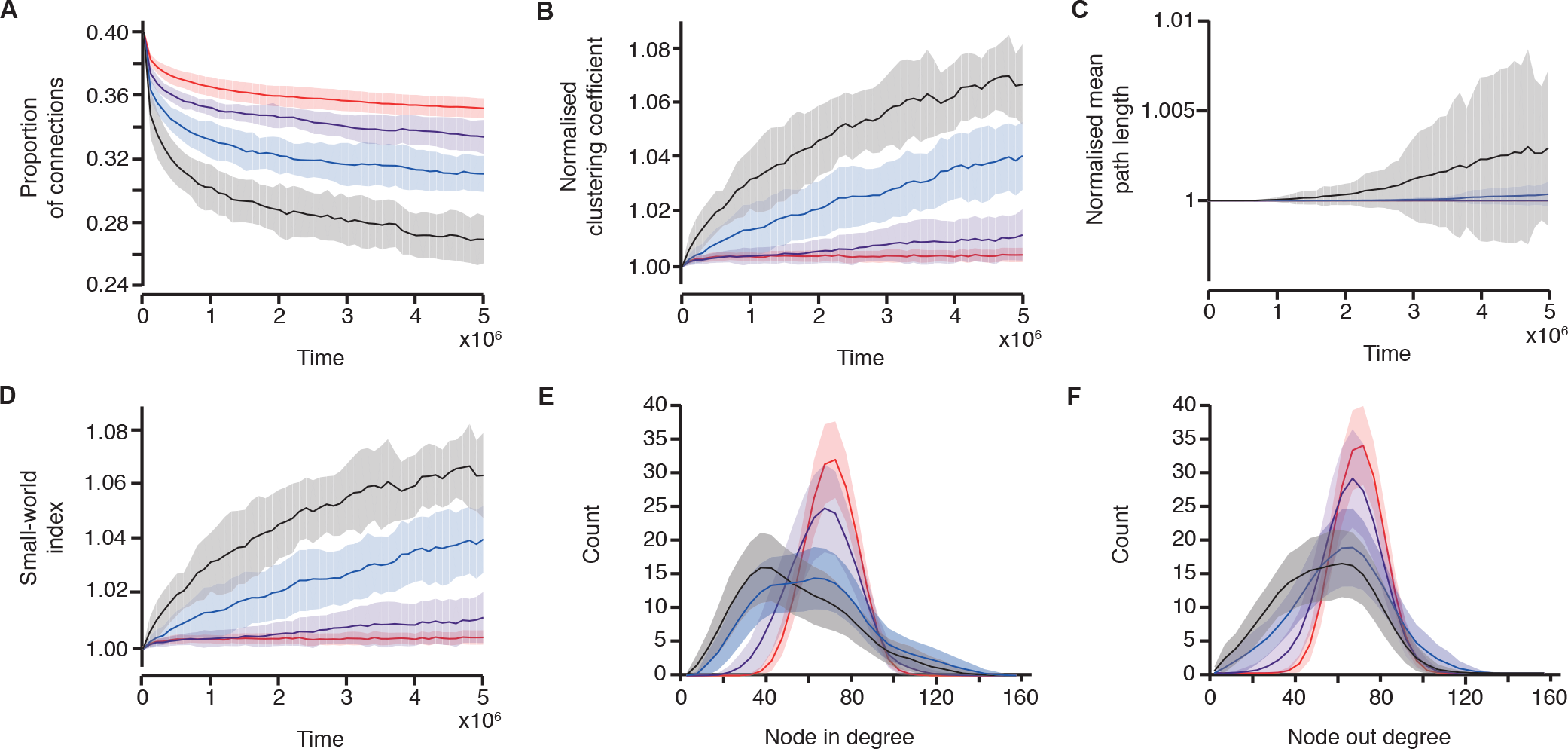
The speed of network evolution is related to the Hurst exponent of the driving input. Changes in network parameters with the networks driven by burst input with temporal correlations with different Hurst exponents: *H* ≈ 0.5 (red), *H* ≈ 0.6 (purple), *H* ≈ 0.7 (blue, same simulations as in Fig. 2), and *H* ≈ 0.8 (black). (A) The proportion of connections in the network, (B) the normalised clustering coefficient, (C) the normalised mean path length (note for *H* ≈ 0.5 and 0.6 the mean path length is equal to one throughout) and (D) the small-world index, across the course of the simulations. (E) The average in degree distributions at the end of the simulations. (F) The average out degree distributions at the end of the simulations. The solid lines indicate the mean across 20 simulations, and the shaded area the standard deviation.

In these simulations, whilst the mean IBI of the external input was equal, the exact distribution of the IBIs was not. However, using the exact same IBI distribution for all simulations, but continuing to vary the Hurst exponent, does not alter the results; the networks driven with LRTCs with higher Hurst exponents exhibit a higher initial rate of change of network parameters (see Fig. S3). This demonstrates that the change in connectivity formation is directly related to the the temporal dynamics of the external input.

### Varying the plasticity parameters

In the simulations to this point, LTD of the likelihood of losing/gaining a connection was set to be slightly stronger than long-term potentiation (LTP) (i.e., *A_D_* = 0.55 and *A_P_* = 0.5, see Methods). This led to the network on average losing connections. We next explored changes in connectivity when these parameters were varied so that potentiation and depression were equal (*A_D_* = *A_P_* = 0.5) and potentiation was greater than depression (*A_D_* = 0.5, *A_P_* = 0.55). With potentiation higher than depression, on average connections are gained, and the normalised clustering coefficient and small-world index increases (Fig. 4). With the potentiation set equal to the level of depression, the change in network parameters across the simulation are very small in comparison, and whilst on average there is also an increase in the proportion of connections, the normalised clustering coefficient and the small-world index, the networks at the end of the simulation do not exhibit a high small-world index. Notably, in both cases the speed of emergence of these properties is higher in the networks driven by bursts which exhibit LRTCs, as was observed with *A_D_* = 0.55 and *A_P_* = 0.5, again demonstrating that the temporal ordering of the burst activity in the external input plays an important role in shaping the network connectivity.

**Figure 4:**
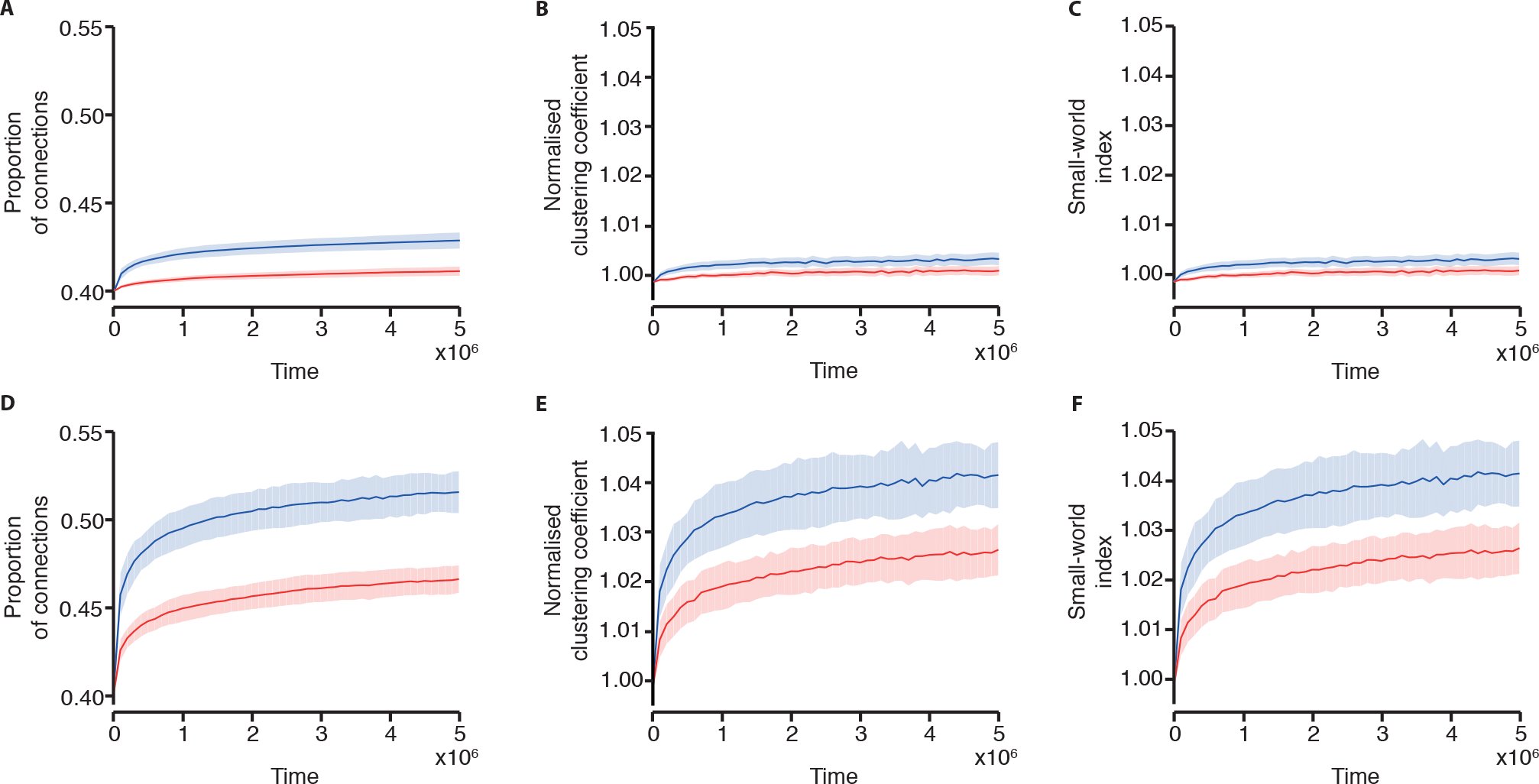
Changes in connectivity are related to the levels of potentiation and depression. (A) The proportion of connections in the network, (B) normalised clustering coefficient, and (C) small-world index across the course of 20 simulations with *A_D_* = *A_P_* = 0.5. (D) The proportion of connections in the network, (E) normalised clustering coefficient, and (F) small-world index across the course of 20 simulations with *A_D_* = 0.5 and *A_P_* = 0.55. The networks are driven with burst dynamics which exhibit LRTCs (*H* ≈ 0.7, blue), compared with the same input randomly shuffled in time (red). Solid lines indicate the mean across the 20 simulations and the shaded area indicates the standard deviation. Results are shown on the same scale for comparison.

We also evaluated changes to the decay constants of the plasticity parameters - *τ*, the decay constant of the spike timing, and *τ_L_*, the decay constant of the likelihood of gaining or losing connections (see Methods). For all previous simulations *τ* = 10 and *τ_L_* = 100. Setting *A_D_* = 0.55 and *A_P_* = 0.5, first we varied *τ* ∈ {5, 6, …, 15}. In all cases the proportion of connections lost, and the normalised clustering coefficient vary according to the Hurst exponent, with the speed of emergence higher in networks which exhibit LRTCs. For lower values of *τ* the overall change in these parameters across the simulation, and the speed of emergence is lower than with *τ* = 10, but small-world properties emerge (see Fig. S4). For higher values, however, when *H* ≈ 0.8 and *H* ≈ 0.7 the number of connections lost at the end of the simulation is high so that the network becomes disconnected. This leads to large variability in the small-world index (Fig. S4) and should be noted as a limitation of our model - eventually it is possible for the networks to become disconnected. A similar pattern also emerges when *τ_L_* is varied with *τ_L_* ∈ {50, 60, …., 150}. For lower values, the speed of emergence is slower than with *τ_L_* = 100 and small-world properties only start to clearly emerge with *H* ≈ 0.8. Thus, it is important to note that with low values of *τ_L_* we do not see a difference in the network properties between the network driven with LRTC and the network driven with random ordering of the bursts in the external input. For higher values of *τ_L_*, with *H* ≈ 0.8, the network becomes disconnected (see Fig. S5), leading to a break down in the small-world properties. However, there is still a distinction in the changes in the proportion of connections and normalised clustering coefficient with the level of the Hurst exponent. Disconnection of the network suggests that high values of both decay constants may lead to pathological network connectivity. This is particularly the case with *H* ≈ 0.8, however, this level of LRTCs is rarely observed in the burst activity of preterm infant EEG (average Hurst exponent estimated using DFA in preterm infants is 0.68 [24]).

### Varying network size and density

Finally, we investigated changes in the network size and density. Previous simulations have all had 200 neurones starting from a random connectivity structure with a density of 40%. Varying the network size did not have a major effect on the network parameters, but reduced the variance see Fig. S6. In contrast, varying the initial connection density had a notable effect on the final network structure, Fig 5. With a low initial connection density of 10%, although there is a difference between the network driven with LRTC and that driven by the external input with random burst ordering in terms of the clustering coefficient, the differences in normalised mean path length between the two lead to both networks having similar smallworld indices throughout the simulations. With a high initial connection density of 50% there is a much slower rate of change of the clustering coefficient and small-world index when the network is driven with LRTCs compared with networks with an initial connection density of 40%. Thus, we importantly note that the difference we observe between networks driven with bursts which exhibit LRTCs and those with random temporal ordering is not always observed but is dependent on the network parameters, including the initial connectivity and the plasticity parameters.

**Figure 5:**
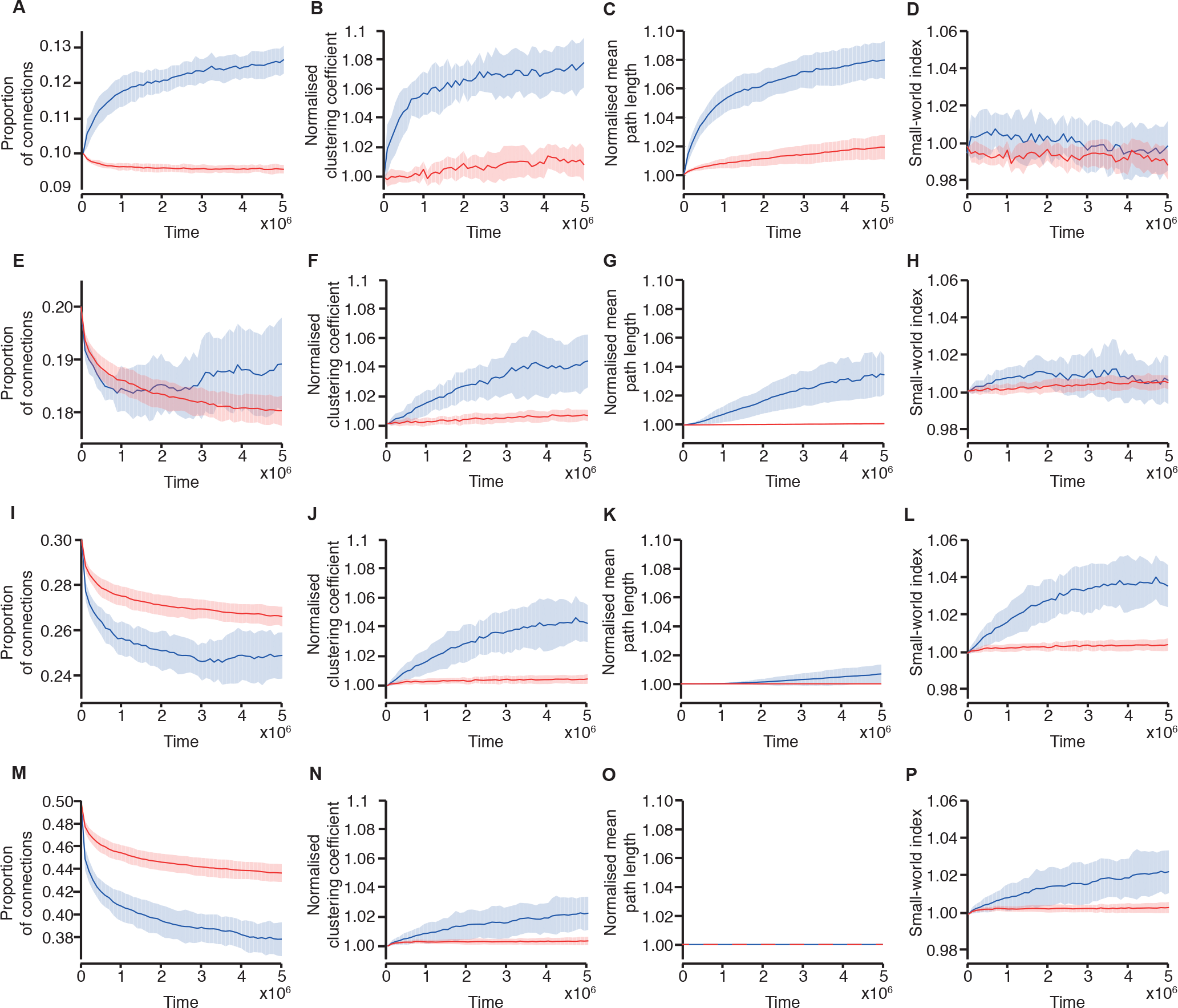
Changes in connectivity in relation to the initial network density. (A,E,I,M) The proportion of connections in the network, (B,F,J,N) normalised clustering coefficient, (C,G,K,O) normalised mean path length and (D,H,L,P) small-world index across the course of 20 simulations with an initial network density of (A-D) 10%, (E-H) 20%, (I-L) 30%, and (M-P) 50%. The networks are driven with burst dynamics which exhibit LRTCs (*H* ≈ 0.7, blue), compared with the same input randomly shuffled in time (red). Solid lines indicate the mean across the 20 simulations and the shaded area indicates the standard deviation.

## Discussion

Here we examined an activity-dependent neuronal network model of connectivity formation in the developing brain. We demonstrate the emergence of small-world topology and the presence of hub neurones, characteristics of brain networks [1, 2, 29, 30]. Furthermore, we found that the temporal ordering of the activity driving the network may be important - when the bursty input driving the network had IBIs which exhibited LRTCs this resulted in faster network evolution compared with when the network was driven by the same input randomly reordered in time (within a range of network parameters). Moreover, network evolution dynamics were related to the magnitude of the Hurst exponent, with a faster speed of emergence of small-world properties in networks driven by external input with higher Hurst exponents.

### Emergence of small-world topology

Starting from networks with initially random connectivity and allowing network connectivity to evolve using a simple Hebbian STDP rule, we observed the emergence of small-world topology and hub neurones. Small-world connectivity has been shown to arise in simple models due to constraints such as efficient neuronal communication and metabolic costs related to neuronal wiring [31], and hub nodes can arise through mechanisms such as preferential attachment [32]. A number of previous authors have also examined more realistic computational models of brain development, including examining axonal growth in molecular gradients [33] and neurite branching [34]. Van Ooyen and Van Pelt showed that a simple model of connectivity formation, with the growth and retraction of circular dendritic and axonal fields based on neuronal activity, results in an equilibrium in connectivity level after an initial overshoot in connectivity consistent with experimental observations [35]. Meisel and Gross further demonstrated that an activity-dependent model of connectivity formation ‘self-organised’ to a balanced connectivity level irrespective of the initial level of connectivity [36]. More recently, Damicelli et al. found that a local Hebbian plasticity rule allowed a network to reorganise to a modular structure [37]. Thus, all of these models demonstrate the importance of neuronal activity in connectivity formation. However, whilst it is known that neuronal activity is essential in the developing brain, both before and after birth [12, 16], we importantly show that the temporal ordering of activity may also play a role in development.

### The role of LRTCs

We find that when driven by burst activity which exhibits LRTCs the network evolves quickly to a state with small-world topology. We speculate that this may be important in development, where quickly transitioning to this type of network will allow for efficient integration and segregation of information in the developing brain. This work was motivated by the observation that inter-burst intervals between bursts of activity in the EEG of preterm infants exhibits LRTCs, with an average estimated Hurst exponent of 0.68 [24]. Evidence suggests that the subplate provides essential input to the cortex during this stage of development [12, 13] so we made the assumption that the burst dynamics of the external input were from a region such as the subplate driving our cortical network. However, the LRTCs in burst dynamics are also passed to the network itself, suggesting that other models which exhibit LRTCs in network activity may also quickly evolve their connectivity. For example, this type of burst dynamics with LRTCs in the inter-burst intervals can emerge in a system driven with continuous external input in the region of a critical state [38]. Although our motivation for this work comes from observations in preterm EEG, LRTCs in adults have been observed at all levels of the nervous system including in spike timing [26]. Moreover, the LRTCs in the burst dynamics of our model are in the external input which we consider to be a population of neurones rather than from single neuronal firing.

### Future directions - understanding pathology in the developing brain

Premature-born infants display altered brain connectivity, indicated by reduced white matter integrity [39, 40] and altered resting state connectivity [41, 42], at term-corrected age [39, 41] and into childhood [40, 42]. Tactile, auditory, visual and noxious stimuli all evoke bursts of activity, observed using EEG, in very preterm infants [8–10]. It is plausible that unexpected sensory exposure, which could disrupt the temporal patterning of the ongoing brain dynamics, in the premature period relates to the long-term neurological problems observed in children who have been born very prematurely [43]. Indeed, the number of painful procedures an infant receives in the premature period is correlated with altered brain development, including lower white matter integrity, and lower cognitive ability at school age [40]. Whether sensory stimuli alter LRTCs in the ongoing EEG of preterm infants is an open question, and a possible extension of this work would be to determine whether sensory input disrupts connectivity formation, which may lead to a better understanding of the long-term effects of premature birth.

Long-term depression of hippocampal synapses in mature cultures has been shown to result in weaker synapses followed by selective elimination of very depressed synapses [44]. Moreover, synaptic pruning shares similar molecular pathways with long-term depression (LTD) [22]. This suggests that connectivity formation in the developing brain may be altered through STDP-like mechanisms, which forms the basis of the model we have used here. In some pathological states, including neurodevelopmental disorders such as autism [45] and schizophrenia [46], brain connectivity is altered, with network architecture which has lower clustering and fewer hub nodes [31]. Both hyper- and hypoconnectivity have been observed in children with autism [47, 48], and LTD dysregulation has been identified in mouse models of autism, leading to the suggestion that alterations in synaptic plasticity and pruning may prevent proper development of brain connectivity [22]. Further exploration of STDP models of connectivity formation, such as the one presented here, may shed light on these disorders.

## Limitations

A significant limitation of this study is the small effect size that was observed. Moreover, the effect was not observed across all parameter choices, for example, with low initial connection density and with low values of the decay constant of the likelihood of gaining or losing connections. Nevertheless, this work demonstrates the possibility that the temporal patterning of external input can lead to differences in structural connectivity formation in neuronal networks and therefore provides motivation for future empirical and experimental investigations to further elucidate the role that the temporal ordering of activity may play in development.

To simplify the model, all neurones within the network were driven by the same external input. Whilst bursts of activity in the preterm EEG known as delta brushes can occur in a diffuse pattern over large cortical areas, this is not always the case, with localised delta brushes also observed [11]. It would not be realistic to model the whole cortex as being driven by one external input, and a useful extension would be to determine how connectivity changes if neurones were to receive different external inputs. As well as the external input, neurones that were connected to each other received input when these neighbouring neurones fired. To avoid either saturation of network firing or quiescence, the weights between individual neurones were evolved according to the level of network connectivity, which can be thought of as a form of homeostatic plasticity [27]. However, a limitation of our approach was that connections could be continually lost (and gained), which can lead to the network forming disconnected components. A possible extension of the work would be to see if maintaining the level of connections within the network, as was done by Damicelli et al. [37] (another form of homeostatic plasticity), but allowing connectivity to continue to evolve under the dynamics of the network, would lead to small-world properties.

## Summary

In conclusion, early spontaneous and sensory driven activity is known to be crucial for the development of connections within neural networks. Here we investigated whether the temporal ordering of burst activity within an external driving input affects connectivity formation in a neuronal network model. Using a STDP model of connectivity formation, we observed that the presence of LRTCs in the ordering of burst activity facilitates the emergence of small-world topology and hub neurones. We suggest that early brain activity, driven by the subplate, leads, through activity-dependent mechanisms, to a small-world cortical network structure, and that LRTCs may play an important role in this connectivity formation warranting further investigation.

## Software availability

All the code for this model can be downloaded here https://github.com/berthouz/BrainDevBursts/tree/1.0 [49].

## Methods

### Neuronal dynamics

Individual neuronal dynamics were modelled as leaky integrate-and-fire neurones described by the differential equation

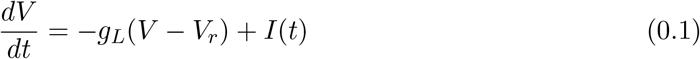

where *V* is the membrane potential of the neurone, *V_r_* is the resting potential, *g_L_* is the leak conductance and *I*(*t*) is the input (both external input and input from other neurones within the system). When the neurone reaches a threshold membrane potential *V_thres_* it fires and is reset to *V_reset_*. For all neurones in the simulations *V_thres_* = −54 *mV*, *V_r_* = −70 *mV*, *V_reset_* = −60 *mV* [50]. The leak conductances were randomly chosen from a normal distribution with mean 0.025 and standard deviation 0.005. This heterogeneity in the conductances leads to heterogeneity in the firing dynamics.

### Neuronal input

The external input to the system was constructed using fractional Gaussian noise; an example of a process that exhibits LRTCs, with a Gaussian data distribution. The IBI sequence was constructed from a random normal distribution with mean ≈ 4.5 and standard deviation ≈ 3. This sequence was ordered according the ordering of the fractional Gaussian noise process, generating LRTCs in the IBI sequence. The external input to the system was constructed from this sequence of IBIs, with bursts with a duration of 5 and an amplitude of 0.8mV between each IBI, see Fig. 1A. We confirmed that the sequence of IBIs constructed in this way exhibited LRTCs indicated by a DFA exponent greater than 0.5 (see *Estimation of the Hurst exponent*). The Hurst exponent of the IBIs was altered by varying the exponent of the fractional Gaussian noise.

We compared the connectivity changes, to connectivity changes within a network evolving under external input with random burst occurrence. This input was constructed by randomly shuffling the IBIs from the original external input. In this way the two inputs are identical in terms of the distribution of the IBIs (and bursts are identical throughout) and it is only the temporal structure of the input that is altered. We also compared the connectivity changes in networks driven with periodic external input. For this case the IBIs were all identical and were set to a duration of 4. Bursts were identical to the external input which exhibited LRTCs, and had a duration of 5 and an amplitude of 0.8*mV*

In Supplementary Figure 1, the rate of input is calculated within moving windows of length 200 by summing the external input within the window and dividing by the window length. As all bursts have equal amplitude, this reflects the temporal ordering of the burst activity within the external input.

The external input was the same to all neurones. Each neurone also received, at each timestep, an input from presynaptic neurones that had fired at the previous time-step. Synaptic weights were equal for all connections and were set to 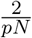 where *N* is the number of neurones in the network and *p* is the proportion of connections in the network. This is calculated at each time step, according to the maximum number of connections which is equal to *N*(*N* − 1) (networks are directed). This update to the synaptic weights can be thought of as a form of homeostatic plasticity - without it the network either stops firing when connectivity falls (the external input alone is not sufficient to make the network fire frequently), or the network starts to fire continually as connectivity is increased. Using this approach, the average levels of activity in the network were maintained across the course of the simulation.

### Estimation of the Hurst exponent

The presence of LRTCs in data can be determined through estimation of the Hurst exponent, *H*. A Hurst exponent of *H* = 0.5 indicates that the data does not exhibit correlations or exhibits short-range correlations only (for example, white noise). A Hurst exponent of 0.5 < *H* < 1 indicates that the data exhibits LRTCs. Here we estimated the Hurst exponent using detrended fluctuation analysis (DFA) [51] which calculates the exponent as the slope of the line of best-fit of the average root-mean-square fluctuations across different box sizes (see Fig. 1C for an example plot and *Peng et al*. [51] for detailed methodology). Briefly, the signal is first integrated and then divided into boxes of equal length, *n*. For each box a least-squares fit to the data is found and the integrated signal is detrended by subtracting this local trend. The root mean square fluctuation, *F*(*n*) is calculated and the process is repeated for different box sizes and the average fluctuation is compared to box size on a double logairthmic plot. The minimum box size was set to 5 IBIs and the maximum to one tenth of the length of the IBI sequence (the recommended maximum window size [52]). This approach has been used by a number of previous authors to determine the presence of LRTCs in data, including neurophysiological data sets [24, 53–58]. As the external input was constructed to have true LRTCs, and these will not be contaminated by noise, it is reasonable to use DFA to calculate a single linear fit to the data. However, in the absence of such prior knowledge, it is more robust to to use maximum likelihood techniques along with model selection methods [59,60].

To examine network firing and periods of activity/inactivity within the network dynamics itself we separated activity using the method of Benayoun et al. [61]. Briefly, two consecutive spikes within a network are separated as distinct bursts if the time difference between them is greater than the average time, *dt*, between consecutive spikes within the total simulation. Thus, a single burst consists of consecutive spikes which are less than *dt* apart. Benayoun et al. used this approach to define avalanches - cascades of network activity - which have periods of separation between them. In this way, avalanches are the same as bursts of activity and so the same approach can be used to determine the bursts here. However, it is worth noting that the term neuronal avalanche is used to define specifically bursts of activity within a network where the distribution of avalanche sizes follows a power-law [62–64].

The average IBI within the network was lower than the average IBI within the external input. This meant that, for the same simulation length, there are more IBIs within the network dynamics, so when calculating the DFA exponents (Fig. 1) the maximum box size is larger in the case of the network firing dynamics than for the external input.

### Connectivity formation

All networks were initially randomly connected with 40 % connectivity, apart from in section *Varying network size and density* where this initial connection density was varied. All networks were directed, and connections were also formed and lost in a direction dependent manner. Connections were updated depending on activity within the network, comparing the firing times between all neuronal pairs. Let *L* be a matrix of values where *L*(*i*, *j*) indicates the likelihood of losing/gaining a connection from neurone *i* (presynaptic) to neurone *j* (postsynaptic). *L*(*i*, *j*) was modified by

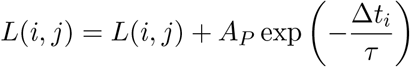

following a spike in neurone *j*, where Δ*t_i_* is the time since the last spike in neurone *i*, *τ* is a decay constant and *A_P_* > 0 is the amplitude change when Δ*t_i_* = 0, and by

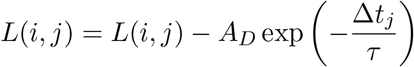

following a spike in neurone *i*, where Δ*t_j_* is the time since the last spike in neurone *j* and *A_D_* > 0 is the amplitude change when Δ*t_j_* = 0.

A connection from *i* to *j* was gained (immediately, if there was not already a connection present) when *L*(*i*, *j*) increased beyond the threshold value *g* = 2. A connection from *i* to *j* was lost when *L*(*i*, *j*) decreased beyond the threshold value *l* = −2. In order to better take into account temporal dynamics within the system (for example if two neurones only spike together rarely) the values of *L* decayed with rate *τ_L_*. Thus, at each time-step:

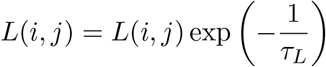

The loss-likelihood was initially set to zero for all connections and as in *Song et al*. [28] depression in initial simulations was set to be slightly stronger than potentiation with *A_P_* = 0.5, *A_D_* = 0.55. We also set *τ* = 10 and *τ_L_* = 100, with relatively slow decay of the likelihood values L allowing for the temporal dynamics of a number of spikes to be taken into account, and the decay of the spike timing *τ* relatively fast so as to allow the temporal dynamics of the external input to take effect (the periods within the external input are relatively short and so to have an effect the ‘memory’ within the system must be of a similar level). We consider changes in the results when these parameters are varied in the final section of the Results.

### Analysis

A number of measures were used to assess how the connection topology changed over the course of the simulations. Firstly, the proportion of connections in the network (the number of connections divided by the number of all possible connections within the network) was used as a straight forward measure to compare whether there were differences in the evolution of the network with different types of the external input. To analyse network topological properties we examined the mean path length and the clustering coefficient of the network. Given any two neurones (or, more generally, nodes within a graph) the shortest path length is the shortest distance needed to be traversed to pass from one node to the other. As we set all the synaptic weights as equal, the shortest path length is equivalent to the lowest number of connections between two neurones. The mean path length is then calculated as the average path length for all pairs of neurones within the network [65]. Given two neurones both connected to a third neurone, the clustering coefficient indicates the likelihood that these two neurones are themselves connected [65]. A random network has a low mean path length and a low clustering coefficient [66]. A number of studies have shown that the neural networks have a similar mean path length to a random network (of the same size and density) but are much more clustered indicating that the brain is a small-world network [2, 29, 31, 66, 67]. Mean path length and clustering coefficients of the networks were calculated using the Brain Connectivity Toolbox, using the functions for binary directed networks [65].

The values of the mean path length, *L*, and clustering coefficient, *C*, are only really meaningful when compared to the average values of random [68, 69] or regular [66] networks of the same size (number of connections and number of neurones). We therefore calculated the values for a random network (*C_rand_* and *L_rand_*) by averaging over the values for 50 random directed networks of the same size. Random networks were constructed in this way for comparison every 100,000 simulation steps (at the same points as the network clustering coefficient and mean path length were calculated). The clustering coefficient and mean path length of a network from the simulations were then normalised by dividing by the average values from the random networks to obtain the normalised clustering coefficient and normalised mean path length respectively. From these values the small-world index can be calculated [68]:

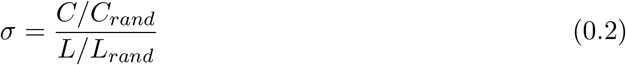

A value of *σ* = 1 indicates that the network is random, whereas *σ* > 1 indicates that the network has small-world properties.

We also compared network degree between the simulations by calculating the degree of the neurones, defined as the number of connections that a neurone makes. As the networks were directed we compared the in-degree distribution (the number of presynaptic neurones a neurone has) and the out-degree distribution (the number of postsynaptic neurones a neurone has) separately.

## Acknowledgments

CH was funded through CoMPLEX (Centre for Mathematics and Physics in the Life Sciences and Experimental Biology), University College London for this project. CH is currently funded by a Wellcome Trust/Royal Society Sir Henry Dale Fellowship. SF acknowledges funding support from the UCLH (University College London Hospitals) Biomedical Research Centre.

## References

[1] Sporns O, Tononi G, Edelman GM. Theoretical neuroanatomy: relating anatomical and functional connectivity in graphs and cortical connection matrices. Cereb Cortex. 2000;10(2):127–41.

[2] van den Heuvel MP, Sporns O. Rich-club organization of the human connectome. J Neurosci. 2011;31(44):15775–86. doi:10.1523/JNEUROSCI.3539-11.2011.

[3] van den Heuvel MP, Kersbergen KJ, de Reus MA, Keunen K, Kahn RS, Groenendaal F, et al. The Neonatal Connectome During Preterm Brain Development. Cereb Cortex. 2015;25(9):3000–13. doi:10.1093/cercor/bhu095.

[4] Ball G, Aljabar P, Zebari S, Tusor N, Arichi T, Merchant N, et al. Rich-club organization of the newborn human brain. Proc Natl Acad Sci U S A. 2014;111(20):7456–61. doi:10.1073/pnas.1324118111.

[5] Kostović I, Jovanov-Milosević N. The development of cerebral connections during the first 20-45 weeks’ gestation. Semin Fetal Neonatal Med. 2006;11(6):415–22. doi:10.1016/j.siny.2006.07.001.

[6] Huttenlocher PR, Dabholkar AS. Regional differences in synaptogenesis in human cerebral cortex. J Comp Neurol. 1997;387(2):167–78.

[7] André M, Lamblin MD, d’Allest AM, Curzi-Dascalova L, Moussalli-Salefranque F, S Nguyen The T, et al. Electroencephalography in premature and full-term infants. Developmental features and glossary. Neurophysiol Clin. 2010;40(2):59–124. doi:10.1016/j.neucli.2010.02.002.

[8] Colonnese MT, Kaminska A, Minlebaev M, Milh M, Bloem B, Lescure S, et al. A conserved switch in sensory processing prepares developing neocortex for vision. Neuron. 2010;67(3):480–98. doi:10.1016/j.neuron.2010.07.015.

[9] Chipaux M, Colonnese MT, Mauguen A, Fellous L, Mokhtari M, Lezcano O, et al. Auditory stimuli mimicking ambient sounds drive temporal delta-brushes in premature infants. PLoS One. 2013;8(11):e79028. doi:10.1371/journal.pone.0079028.

[10] Fabrizi L, Slater R, Worley A, Meek J, Boyd S, Olhede S, et al. A shift in sensory processing that enables the developing human brain to discriminate touch from pain. Curr Biol. 2011;21(18):1552–8. doi:10.1016/j.cub.2011.08.010.

[11] Milh M, Kaminska A, Huon C, Lapillonne A, Ben-Ari Y, Khazipov R. Rapid cortical oscillations and early motor activity in premature human neonate. Cereb Cortex. 2007;17(7):1582–94. doi:10.1093/cercor/bhl069.

[12] Dupont E, Hanganu IL, Kilb W, Hirsch S, Luhmann HJ. Rapid developmental switch in the mechanisms driving early cortical columnar networks. Nature. 2006;439(7072):79–83. doi:10.1038/nature04264.

[13] Vanhatalo S, Lauronen L. Neonatal SEP - back to bedside with basic science. Semin Fetal Neonatal Med. 2006;11(6):464–70. doi:10.1016/j.siny.2006.07.009.

[14] Kanold PO. Subplate neurons: crucial regulators of cortical development and plasticity. Front Neuroanat. 2009;3:16. doi:10.3389/neuro.05.016.2009.

[15] Arichi T, Whitehead K, Barone G, Pressler R, Padormo F, Edwards AD, et al. Localization of spontaneous bursting neuronal activity in the preterm human brain with simultaneous EEG-fMRI. Elife. 2017;6. doi:10.7554/eLife.27814.

[16] Katz LC, Shatz CJ. Synaptic activity and the construction of cortical circuits. Science. 1996;274(5290):1133–8.

[17] Tolner EA, Sheikh A, Yukin AY, Kaila K, Kanold PO. Subplate neurons promote spindle bursts and thalamocortical patterning in the neonatal rat somatosensory cortex. J Neurosci. 2012;32(2):692–702. doi:10.1523/JNEUROSCI.1538-11.2012.

[18] Ghosh A, Shatz CJ. Involvement of subplate neurons in the formation of ocular dominance columns. Science. 1992;255(5050):1441–3.

[19] Kanold PO, Kara P, Reid RC, Shatz CJ. Role of subplate neurons in functional maturation of visual cortical columns. Science. 2003;301(5632):521–5. doi:10.1126/science.1084152.

[20] Schmidt JT, Eisele LE. Stroboscopic illumination and dark rearing block the sharpening of the regenerated retinotectal map in goldfish. Neuroscience. 1985;14(2):535–46.

[21] Weliky M, Katz LC. Disruption of orientation tuning in visual cortex by artificially correlated neuronal activity. Nature. 1997;386(6626):680–5. doi:10.1038/386680a0.

[22] Piochon C, Kano M, Hansel C. LTD-like molecular pathways in developmental synaptic pruning. Nat Neurosci. 2016;19(10):1299–310. doi:10.1038/nn.4389.

[23] Iyer KK, Roberts JA, Hellström-Westas L, Wikström S, Hansen Pupp I, Ley D, et al. Cortical burst dynamics predict clinical outcome early in extremely preterm infants. Brain. 2015;138(Pt 8):2206–18. doi:10.1093/brain/awv129.

[24] Hartley C, Berthouze L, Mathieson SR, Boylan GB, Rennie JM, Marlow N, et al. Longrange temporal correlations in the EEG bursts of human preterm babies. PLoS One. 2012;7(2):e31543. doi:10.1371/journal.pone.0031543.

[25] Ben-Ari Y. Excitatory actions of GABA during development: the nature of the nurture. Nat Rev Neurosci. 2002;3(9):728–39. doi:10.1038/nrn920.

[26] Bhattacharya J, Edwards J, Mamelak AN, Schuman EM. Long-range temporal correlations in the spontaneous spiking of neurons in the hippocampal-amygdala complex of humans. Neuroscience. 2005;131(2):547–55. doi:10.1016/j.neuroscience.2004.11.013.

[27] Turrigiano G. Too many cooks? Intrinsic and synaptic homeostatic mechanisms in cortical circuit refinement. Annu Rev Neurosci. 2011;34:89–103. doi:10.1146/annurevneuro-060909-153238.

[28] Song S, Miller KD, Abbott LF. Competitive Hebbian learning through spike-timingdependent synaptic plasticity. Nat Neurosci. 2000;3(9):919–26. doi:10.1038/78829.

[29] Bassett DS, Bullmore E. Small-world brain networks. Neuroscientist. 2006;12(6):512–23. doi:10.1177/1073858406293182.

[30] Sporns O, Chialvo DR, Kaiser M, Hilgetag CC. Organization, development and function of complex brain networks. Trends Cogn Sci. 2004;8(9):418–25. doi:10.1016/j.tics.2004.07.008.

[31] Sporns O. The non-random brain: efficiency, economy, and complex dynamics. Front Comput Neurosci. 2011;5:5. doi:10.3389/fncom.2011.00005.

[32] Barabasi, Albert. Emergence of scaling in random networks. Science. 1999;286(5439):509–12.

[33] Goodhill GJ, Gu M, Urbach JS. Predicting axonal response to molecular gradients with a computational model of filopodial dynamics. Neural Comput. 2004;16(11):2221–43. doi:10.1162/0899766041941934.

[34] Kiddie G, McLean D, Van Ooyen A, Graham B. Biologically plausible models of neurite outgrowth. Prog Brain Res. 2005;147:67–80. doi:10.1016/S0079-6123(04)47006-X.

[35] Van Ooyen A, Van Pelt J. Activity-dependent outgrowth of neurons and overshoot phenoman in developing neural networks. J Theor Biol. 1994;167:27–43. doi:10.1016/S0079-6123(08)60544-0.

[36] Meisel C, Gross T. Adaptive self-organization in a realistic neural network model. Physical Review E. 2009;80(6):061917. doi:DOI 10.1103/PhysRevE.80.061917.

[37] Damicelli F, Hilgetag CC, Hütt MT, Messé A. Modular topology emerges from plasticity in a minimalistic excitable network model. Chaos. 2017;27(4):047406. doi:10.1063/1.4979561.

[38] Hartley C, Taylor TJ, Kiss IZ, Farmer SF, Berthouze L. Identification of Criticality in Neuronal Avalanches: II. A Theoretical and Empirical Investigation of the Driven Case. J Math Neurosci. 2014;4:9. doi:10.1186/2190-8567-4-9.

[39] Rose SE, Hatzigeorgiou X, Strudwick MW, Durbridge G, Davies PSW, Colditz PB. Altered white matter diffusion anisotropy in normal and preterm infants at term-equivalent age. Magn Reson Med. 2008;60(4):761–7. doi:10.1002/mrm.21689.

[40] Vinall J, Miller SP, Bjornson BH, Fitzpatrick KPV, Poskitt KJ, Brant R, et al. Invasive procedures in preterm children: brain and cognitive development at school age. Pediatrics. 2014;133(3):412–21. doi:10.1542/peds.2013-1863.

[41] Smyser CD, Snyder AZ, Shimony JS, Mitra A, Inder TE, Neil JJ. Resting-State Network Complexity and Magnitude Are Reduced in Prematurely Born Infants. Cereb Cortex. 2016;26(1):322–333. doi:10.1093/cercor/bhu251.

[42] Damaraju E, Phillips JR, Lowe JR, Ohls R, Calhoun VD, Caprihan A. Resting-state functional connectivity differences in premature children. Front Syst Neurosci. 2010;4. doi:10.3389/fnsys.2010.00023.

[43] Grunau RE. Neonatal pain in very preterm infants: long-term effects on brain, neurodevelopment and pain reactivity. Rambam Maimonides Med J. 2013;4(4):e0025. doi:10.5041/RMMJ.10132.

[44] Wiegert JS, Oertner TG. Long-term depression triggers the selective elimination of weakly integrated synapses. Proc Natl Acad Sci U S A. 2013;110(47):E4510–9. doi:10.1073/pnas.1315926110.

[45] Just MA, Cherkassky VL, Keller TA, Kana RK, Minshew NJ. Functional and anatomical cortical underconnectivity in autism: evidence from an FMRI study of an executive function task and corpus callosum morphometry. Cereb Cortex. 2007;17(4):951–61. doi:10.1093/cercor/bhl006.

[46] Rubinov M, Knock SA, Stam CJ, Micheloyannis S, Harris AWF, Williams LM, et al. Small-world properties of nonlinear brain activity in schizophrenia. Hum Brain Mapp. 2009;30(2):403–16. doi:10.1002/hbm.20517.

[47] Supekar K, Uddin LQ, Khouzam A, Phillips J, Gaillard WD, Kenworthy LE, et al. Brain hyperconnectivity in children with autism and its links to social deficits. Cell Rep. 2013;5(3):738–47. doi:10.1016/j.celrep.2013.10.001.

[48] Mostofsky SH, Powell SK, Simmonds DJ, Goldberg MC, Caffo B, Pekar JJ. Decreased connectivity and cerebellar activity in autism during motor task performance. Brain. 2009;132(Pt 9):2413–25. doi:10.1093/brain/awp088.

[49] Hartley C, Berthouze L. Code for Temporal ordering of input modulates connectivity formation in a developmental neuronal network model of the cortex, doi:10.5281/zenodo.1297410;doi:10.5281/zenodo.1297410.

[50] Liu YH, Wang XJ. Spike-frequency adaptation of a generalized leaky integrate-and-fire model neuron. J Comput Neurosci. 2001;10(1):25–45.

[51] Peng CK, Havlin S, Stanley HE, Goldberger AL. Quantification of scaling exponents and crossover phenomena in nonstationary heartbeat time series. Chaos. 1995;5(1):82–7. doi:10.1063/1.166141.

[52] Hu K, Ivanov PC, Chen Z, Carpena P, Stanley HE. Effect of trends on detrended fluctuation analysis. Phys Rev E Stat Nonlin Soft Matter Phys. 2001;64(1 Pt 1):011114. doi:10.1103/PhysRevE.64.011114.

[53] Linkenkaer-Hansen K, Nikouline VV, Palva JM, Ilmoniemi RJ. Long-range temporal correlations and scaling behavior in human brain oscillations. J Neurosci. 2001;21(4):1370–7.

[54] Linkenkaer-Hansen K, Nikulin VV, Palva JM, Kaila K, Ilmoniemi RJ. Stimulus-induced change in long-range temporal correlations and scaling behaviour of sensorimotor oscillations. Eur J Neurosci. 2004;19(1):203–11.

[55] Nikulin VV, Brismar T. Long-range temporal correlations in alpha and beta oscillations: effect of arousal level and test-retest reliability. Clin Neurophysiol. 2004;115(8):1896–908. doi:10.1016/j.clinph.2004.03.019.

[56] Nikulin VV, Brismar T. Long-range temporal correlations in electroencephalographic oscillations: Relation to topography, frequency band, age and gender. Neuroscience. 2005;130(2):549–58. doi:10.1016/j.neuroscience.2004.10.007.

[57] Berthouze L, James LM, Farmer SF. Human EEG shows long-range temporal correlations of oscillation amplitude in Theta, Alpha and Beta bands across a wide age range. Clin Neurophysiol. 2010;121(8):1187–97. doi:10.1016/j.clinph.2010.02.163.

[58] Smit DJA, de Geus EJC, van de Nieuwenhuijzen ME, van Beijsterveldt CEM, van Baal GCM, Mansvelder HD, et al. Scale-free modulation of resting-state neuronal oscillations reflects prolonged brain maturation in humans. J Neurosci. 2011;31(37):13128–36. doi:10.1523/JNEUROSCI.1678-11.2011.

[59] Botcharova M, Farmer SF, Berthouze L. A maximum likelihood based technique for validating detrended fluctuation analysis (ML-DFA). arXiv. 2013; p. 1306.5075.

[60] Ton R, Daffertshofer A. Model selection for identifying power-law scaling. Neuroimage. 2016;136:215–26. doi:10.1016/j.neuroimage.2016.01.008.

[61] Benayoun M, Cowan JD, van Drongelen W, Wallace E. Avalanches in a stochastic model of spiking neurons. PLoS Comput Biol. 2010;6(7):e1000846. doi:10.1371/journal.pcbi.1000846.

[62] Beggs JM, Plenz D. Neuronal avalanches in neocortical circuits. J Neurosci. 2003;23(35):11167–77.

[63] Beggs JM, Plenz D. Neuronal avalanches are diverse and precise activity patterns that are stable for many hours in cortical slice cultures. J Neurosci. 2004;24(22):5216–29. doi:10.1523/JNEUROSCI.0540-04.2004.

[64] Taylor TJ, Hartley C, Simon PL, Kiss IZ, Berthouze L. Identification of Criticality in Neuronal Avalanches: I. A Theoretical Investigation of the Non-driven Case. J Math Neurosci. 2013;3(1):5. doi:10.1186/2190-8567-3-5.

[65] Rubinov M, Sporns O. Complex network measures of brain connectivity: uses and interpretations. Neuroimage. 2010;52(3):1059–69. doi:10.1016/j.neuroimage.2009.10.003.

[66] Watts DJ, Strogatz SH. Collective dynamics of ‘small-world’ networks. Nature. 1998;393(6684):440–2. doi:10.1038/30918.

[67] Bassett DS, Meyer-Lindenberg A, Achard S, Duke T, Bullmore E. Adaptive reconfiguration of fractal small-world human brain functional networks. Proc Natl Acad Sci U S A. 2006;103(51):19518–23. doi:10.1073/pnas.0606005103.

[68] Humphries MD, Gurney K. Network ‘small-world-ness’: a quantitative method for determining canonical network equivalence. PLoS One. 2008;3(4):e0002051. doi:10.1371/journal.pone.0002051.

[69] Gritsun TA, le Feber J, Rutten WLC. Growth dynamics explain the development of spatiotemporal burst activity of young cultured neuronal networks in detail. PLoS One. 2012;7(9):e43352. doi:10.1371/journal.pone.0043352.

