## Supplementary Material for "Temporal ordering of input modulates connectivity formation in a developmental neuronal network model of the cortex"

**Fig. S1 The initial rate of input to the network is similar whether the external input sequence of IBIs exhibits LRTC or is shuffled.** The average rate of input at the start of simulations when the external input exhibits LRTCs (blue) compared with the input randomly shuffled in time (red). This is shown for (A) the first 2% and (B) the first 0.2% of the simulations. Solid lines indicate the mean rate of input across 20 simulations, and the shaded area indicates the standard deviations.

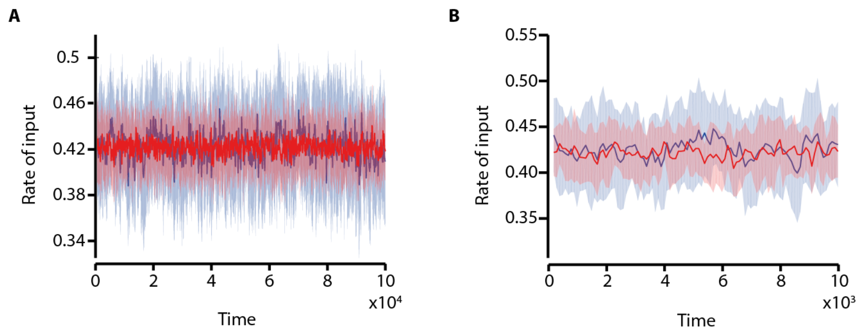

**Fig. S2 The speed of emergence of small-world properties at the start of simulations is not related to the average rate of input at the start of the simulations.** (A,B,C) Moving average of the first 1000 IBIs (averaging across 100) for example IBI sequences which exhibit LRTCs (blue, $H \approx 0.7$) and are randomly shuffled (red, $H \approx 0.5$). (A,B) An example of a IBI sequence which exhibits LRTCs compared with a shuffled sequence. (C) The two IBI sequences which exhibit LRTCs are compared. Note that the full simulations have over 500,000 IBIs so the sequences in these figures equate to approximately the first 0.2% of the simulation. (D,E,F) The proportion of connections in the network across the full simulations for the equivalent IBI sequences shown in the row above. The initial rate of input to the network, reflected in the IBI sequence, does not affect the speed at which the network evolves.

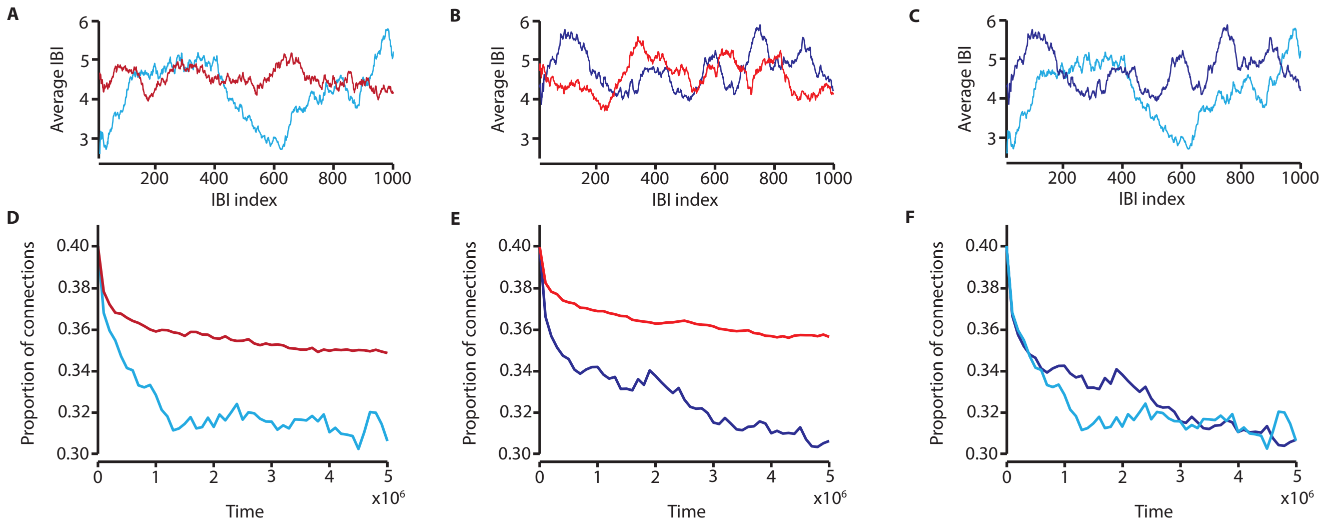

**Fig. S3 The rate of network evolution is related to the Hurst exponent of the driving input independent of the overall IBI distribution.** Changes in network parameters with the networks driven by burst input with identical IBI distributions (across all simulations in this figure) but with temporal correlations with different Hurst exponents: $H \approx 0.5$ (red), $H \approx 0.6$ (purple), $H \approx 0.7$ (blue), and $H \approx 0.8$ (black). (A) The proportion of connections in the network, (B) the normalised clustering coefficient, (C) the normalised mean path length (note for $H \approx 0.5$ and 0.6 the mean path length is equal to one throughout) and (D) the small-world index across the course of the simulations. (E) The average in degree distributions at the end of the simulations. The solid lines indicate the mean across 20 simulations, and the shaded area the standard deviation. Fig. 3 demonstrated that the speed of emergence of small-world properties is dependent on the magnitude of the Hurst exponent. However, the IBI distribution in these simulations was not identical as is the case here. This confirms that the differences in the evolution of network parameters is related to the magnitude of the Hurst exponent rather than the IBI distribution.

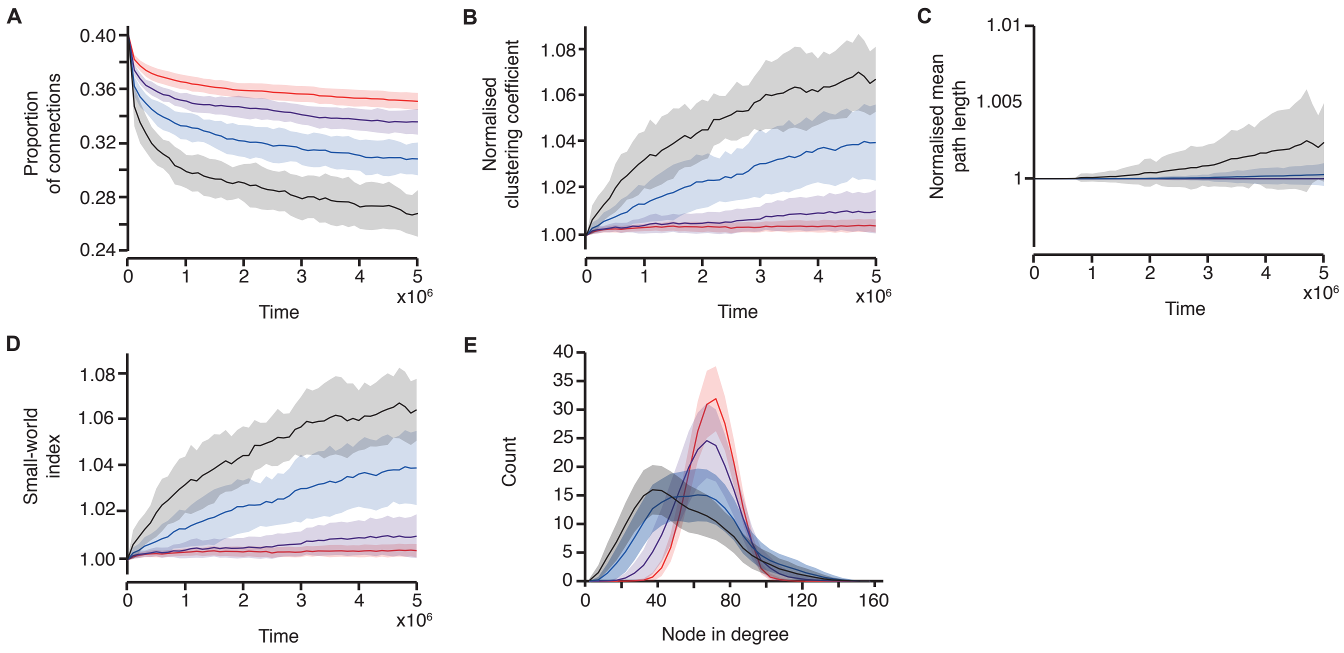

**Fig. S4 Network evolution varies with the decay constant of spike timing.** Changes in the network parameters with different values of the decay constant of spike timing, $\tau$. $\tau$ was varied between (A,B,C,D) $\tau = 5$ and (E,F,G,H) $\tau = 15$. For lower values the network changes are small but small-world properties emerge, and the rate of this emergence is dependent on the Hurst exponent. For higher values of $\tau$ with $H \approx 0.7, 0.8$ the network becomes disconnected leading to a break down of the small-world properties. (A, E) The proportion of connections in the network, (B, F) the normalised clustering coefficient, (C, G) the small-world index, and (D, H) the number of components across the course of simulations with $H \approx 0.5$ (red), $H \approx 0.6$ (purple), $H \approx 0.7$ (blue), and $H \approx 0.8$ (black). Note that in (D) the number of components is equal to one throughout all simulations. Solid lines indicate the mean across 20 simulations, and the shaded area the standard deviation.

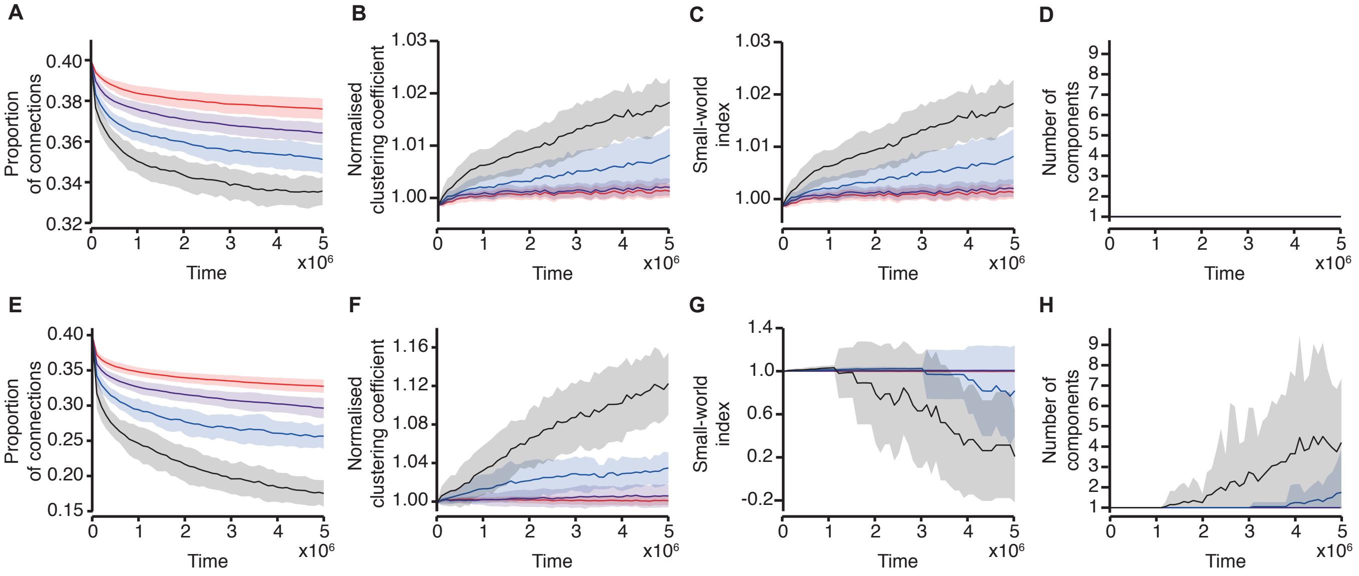

**Fig. S5 Network evolution varies with the decay constant of the likelihood of gaining or losing connections.** Changes in the network parameters with different values of the decay constant $\tau_L$. $\tau_L$ was varied between (A,B,C,D) $\tau_L = 50$ and (E,F,G,H) $\tau_L = 150$. For low values of $\tau_L$ the network changes are very small but small-world properties start to emerge with $H \approx 0.8$. For higher values of $\tau$ with $H \approx 0.7, 0.8$ the network starts to become disconnected during the course of the simulations. However, there is still a clear distinction in the changes in the proportion of connections and normalised clustering coefficient with different values of the Hurst exponent. (A, E) The proportion of connections in the network, (B, F) the normalised clustering coefficient, (C, G) the small-world index, and (D, H) the number of components across the course of simulations with $H \approx 0.5$ (red), $H \approx 0.6$ (purple), $H \approx 0.7$ (blue), and $H \approx 0.8$ (black). Note that in (D) the number of components is equal to one throughout all simulations. Solid lines indicate the mean across 20 simulations, and the shaded area the standard deviation.

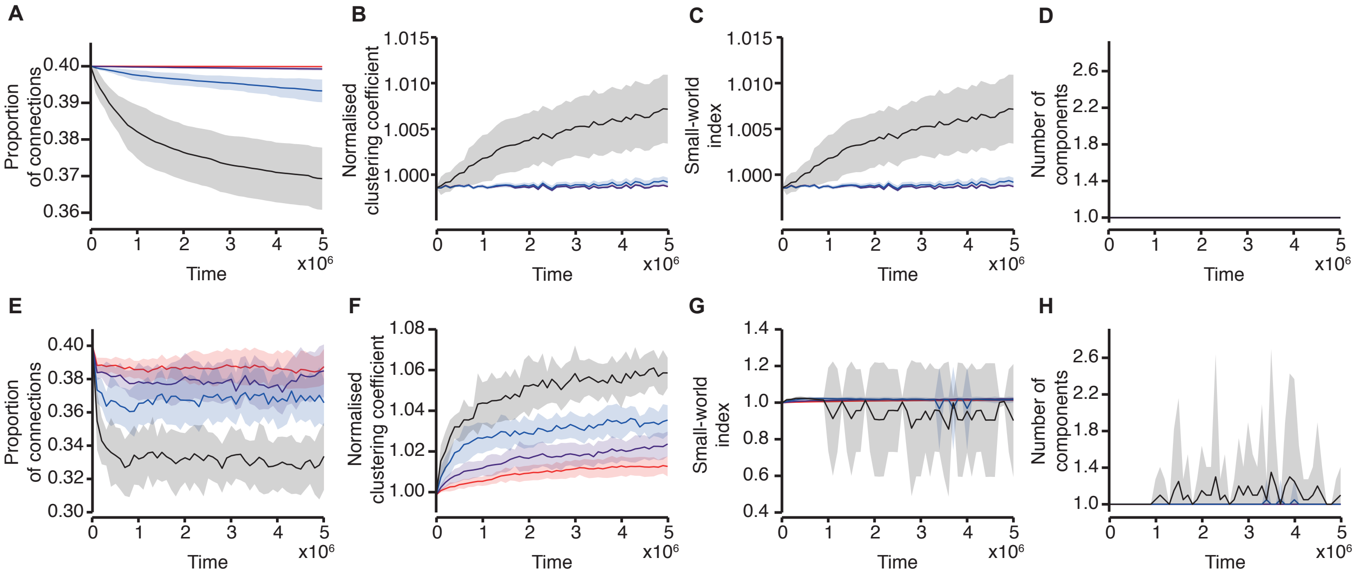

**Fig. S6 Changes in connectivity in relation to the size of the network.** (A,E,I,M) The proportion of connections in the network, (B,F,J,N) normalised clustering coefficient, (C,G,K,O) normalised mean path length and (D,H,L,P) small-world index across the course of 20 simulations with a network size of (A-D) N=100, (E-H) N=500, (I-L) N=1000, and (M-P) N=2000. The networks are driven with burst dynamics which exhibit LRTCs ($H \approx 0.7$, blue), compared with the same input randomly shuffled in time (red). Solid lines indicate the mean across the 20 simulations and the shaded area indicates the standard deviation. Results are shown on the same scale for comparison.

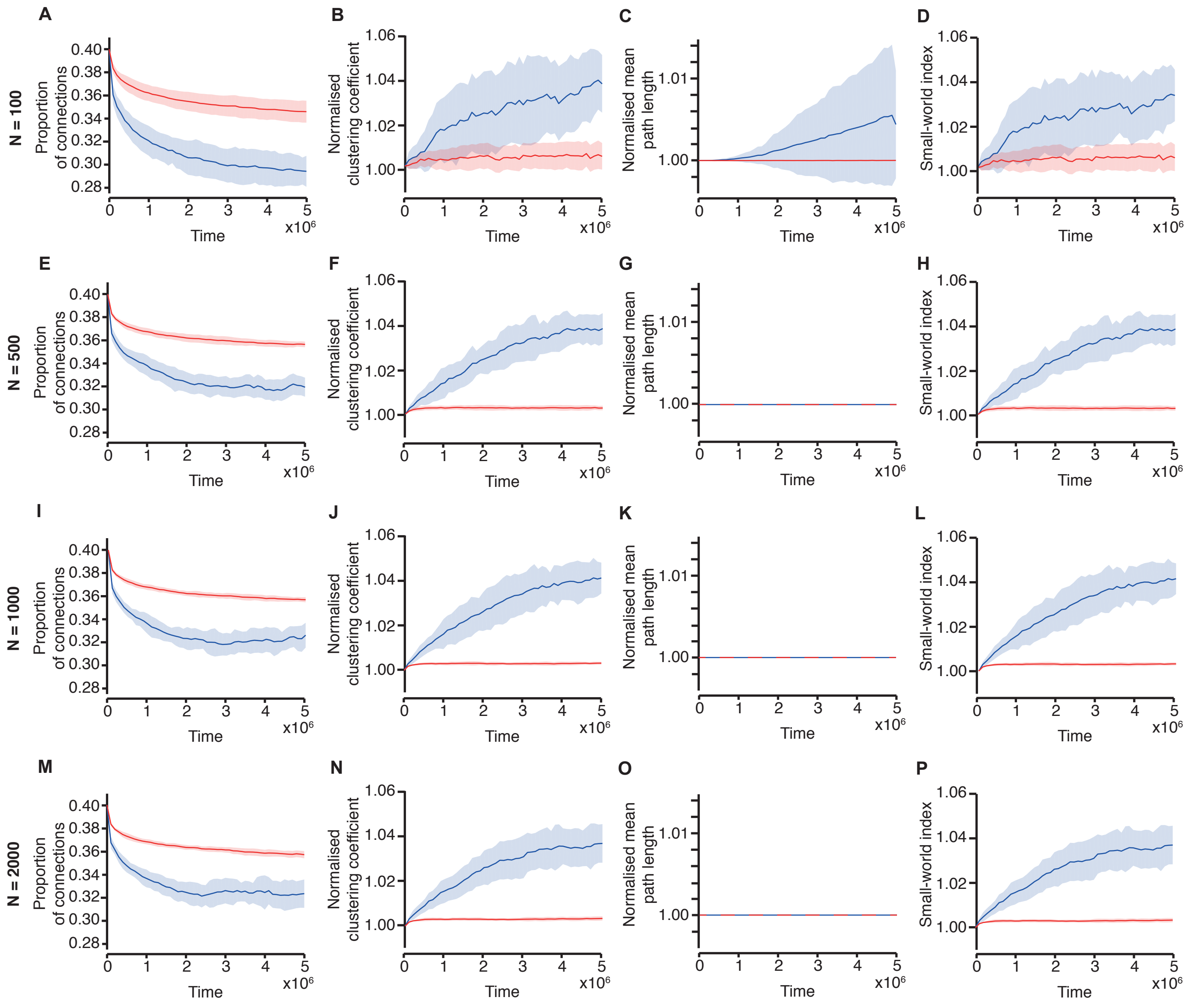
